# The Role of Ebola Virus VP24 Nuclear Trafficking Signals in Infectious Particle Production

**DOI:** 10.1101/2024.03.13.584761

**Authors:** Olivia A. Vogel, Elias Nafziger, Anurag Sharma, H. Amalia Pasolli, Robert A. Davey, Christopher F. Basler

## Abstract

Ebola virus (EBOV) protein VP24 carries out at least two critical functions. It promotes condensation of viral nucleocapsids, which is crucial for infectious virus production, and it suppresses interferon (IFN) signaling, which requires interaction with the NPI-1 subfamily of importin-α (IMPA) nuclear transport proteins. Interestingly, over-expressed IMPA leads to VP24 nuclear accumulation and a carboxy-terminus nuclear export signal (NES) has been reported, suggesting that VP24 may undergo nuclear trafficking. For the first time, we demonstrate that NPI-1 IMPA overexpression leads to the nuclear accumulation of VP24 during EBOV infection. To assess the functional impact of nuclear trafficking, we generated tetracistronic minigenomes encoding VP24 nuclear import and/or export signal mutants. The minigenomes, which also encode *Renilla* luciferase and viral proteins VP40 and GP, were used to generate transcription and replication competent virus-like particles (trVLPs) that can be used to assess EBOV RNA synthesis, gene expression, entry and viral particle production. With this system, we confirmed that NES or IMPA binding site mutations altered VP24 nuclear localization, demonstrating functional trafficking signals. While these mutations minimally affected transcription and replication, the trVLPs exhibited impaired infectivity and formation of shortened nucleocapsids for the IMPA binding mutant. For the NES mutants, infectivity was reduced approximately 1000-fold. The NES mutant could still suppress IFN signaling but failed to promote nucleocapsid formation. To determine whether VP24 nuclear export is required for infectivity, the residues surrounding the wildtype NES were mutated to alanine or the VP24 NES was replaced with the Protein Kinase A Inhibitor NES. While nuclear export remained intact for these mutants, infectivity was severely impaired. These data demonstrate that VP24 undergoes nuclear trafficking and illuminates a separate and critical role for the NES and surrounding sequences in infectivity and nucleocapsid assembly.

**Author Statement:** Ebola virus (EBOV) viral protein 24 (VP24) interacts with importin-α (IMPA) nuclear transport proteins to inhibit interferon signaling. VP24 also contains a nuclear export signal (NES). The capacity to bind IMPA and the presence of a NES suggest that VP24 traffics to the nucleus. However, this had not been demonstrated in the context of EBOV replication. In addition, whether these signals influence virus replication beyond effects on interferon responses has been unclear. Here, we demonstrate that VP24 traffics to the nucleus via IMPA during EBOV infection and in the context of an EBOV transcription and replication competent virus-like particle (trVLP) assay. Using the trVLP system, we also confirmed that VP24 possesses a functional nuclear export signal (NES). Mutations disrupting VP24-IMPA interaction or nuclear export reduced or abrogated trVLP infectivity, respectively. VP24 is known to be necessary for viral nucleocapsid maturation. The VP24 IMPA interface mutants yielded shortened nucleocapsids while VP24 NES mutations led to loss of nucleocapsid formation. Interestingly, VP24 mutants with intact NES function but altered sequences near the NES remain incapable of producing nucleocapsids. Together, these data demonstrate VP24 undergoes nuclear trafficking and reveals additional roles for both the IMPA interaction interface and the NES in nucleocapsid assembly.

## Introduction

Ebola virus (EBOV) is an enveloped, non-segmented negative strand RNA virus from the family *Filoviridae* [1]. The family includes two genera that contain viruses associated with severe human disease, *Orthoebolavirus* (formerly *Ebolavirus*), which includes EBOV, and *Orthomarburgvirus* (formerly *Marburgvirus*), which includes Marburg virus (MARV)[1]. The family also includes two viruses thus far detected in bats and that are classified in additional genera, Lloviu virus (LLOV) of the genus *Cuevavirus*, and Měnglà virus (MLAV) in the genus *Dianlovirus* [1–4]. Among the filoviruses, EBOV has caused the greatest number of documented outbreaks and the largest and most deadly outbreaks, as exemplified by the 2014-2016 West Africa EBOV outbreak that resulted in more than 28,000 cases and 11,000 deaths [5, 6].

The EBOV genome has seven distinct transcription units each encoding one of seven viral structural proteins called the nucleoprotein (NP), viral protein 35 (VP35), VP40, the glycoprotein (GP), VP30, VP24, and the large (L) protein. Non-structural, secreted products of the GP gene are also produced via transcriptional editing by the L protein [5]. Among the viral proteins, VP24 is notable because it is common to filoviruses but lacks obvious homologues in other viral families. VP24 was originally described as a minor matrix protein based on its presence in viral particles and the apparent capacity of VP24 from EBOV or MARV to associate with cellular membranes [7–9]. However, VP24 lacks measurable capacity to directly bind lipids [10]. Rather, VP24 interacts with NP and VP35 to promote formation of nucleocapsid-like structures (NCLS) nearly identical to the mature nucleocapsids found in virions [11]. The capacity of VP24 to promote NCLS condensation likely accounts for its capacity to inhibit EBOV transcription and replication when overexpressed [12, 13]. Based on cryoelectron tomography studies, mature nucleocapsids have an inner layer of NP, which forms a left-handed helical structure. Extending from the inner helix are “boomerang-shaped” projections that are likely formed by VP35 and VP24 and may cause the nucleocapsids to become more rigid [14–16]. In addition, co-expression of VP24 with NP and VP35 is sufficient to allow actin-dependent, directional transport of NCLS, likely providing a mechanism by which nucleocapsids traffic to sites of viral budding at the plasma membrane [17]. These findings likely explain why VP24 plays an essential role in generation of infectious virus particles [18–21].

Filovirus VP24 proteins also engage host signaling pathways, with VP24s from the *Ebolavirus* and *Cuevavirus* genera counter STAT1-dependent antiviral responses and MARV VP24 interacts with host protein Keap1 to activate cytoprotective anti-oxidant responses [22–30]. There is also evidence for EBOV and LLOV VP24 inhibition of type I and type III interferon (IFN) induction pathways [26, 31–33]. EBOV VP24 blocks cellular responses to IFNs by impeding the nuclear accumulation of tyrosine phosphorylated STAT1 [22–24, 34]. This block is mediated by VP24 interaction with the carboxyterminal region of a subset of importin alpha (IMPA) nuclear transport factors, IMPA5, 6 and 7, known collectively as the NPI-1 subfamily. These are the same IMPAs that mediate nuclear import of tyrosine-phosphorylated, dimerized STAT1 [22–24]. VP24 binds the non-classical nuclear localization signal (NLS) binding site that is used by STAT1 for its nuclear import, accounting for impaired STAT1 nuclear localization following addition of IFN to VP24-expressing or EBOV-infected cells [22–24].

Interaction of VP24 with IMPA proteins suggests a capacity to traffic to the nucleus, although at steady state in infected cells it exhibits a cytoplasmic localization and accumulates at viral inclusion bodies [9, 35]. Transfected VP24 interacts with a number of typically nuclear proteins, and over-expression of IMPA5 promotes nuclear accumulation of transfected VP24, suggesting that VP24 can traffic to the nucleus [36, 37]. Studies on transfected VP24 also identified a CRM1-dependent nuclear export signal (NES) near the C-terminus [38]. However, the functional significance of the NES for EBOV replication has not been assessed. The role of VP24-IMPA interaction in EBOV infection has been studied in more depth. For example, recombinant EBOVs bearing mutations that decrease VP24 affinity for IMPA exhibited growth attenuation in cell culture, and this appeared to be due to IFN-related and IFN-independent mechanisms [39–41]. However, the functional and mechanistic significance of VP24-IMPA interaction beyond suppression of IFN signaling remains incompletely understood.

Here, we sought to determine how the VP24-IMPA interaction interface and the NES contribute to the outcome of infection. EBOV infection studies demonstrate the capacity of NPI-1 subfamily member IMPA7 to direct VP24 into the nucleus. Using the EBOV transcription-replication competent virus-like particle (trVLP) life cycle modeling assay, we demonstrate that VP24 has functional nuclear import and export signals. Mutating the VP24-IMPA interface had a modest impact on infectivity, while mutating the VP24 NES had profound negative effects on infectivity. Through co-immunoprecipitation experiments and reporter assays, we determined that the defect in infectivity observed in the NES mutants was independent of VP24’s role in the inhibition of IFN signaling. Instead, the VP24 NES mutants are defective in nucleocapsid assembly, abrogating infectivity. We also demonstrated that VP24 nuclear trafficking is not sufficient for nucleocapsid assembly; however, the NES and surrounding residues are important for proper nucleocapsid assembly and infectivity. These results indicate that both the VP24-IMPA interface and VP24 NES serve multiple purposes in the EBOV life cycle with roles in regulating nuclear trafficking, inhibition of IFN signaling, and nucleocapsid assembly.

## Results

### VP24 Nuclear Trafficking Signals are Important for trVLP Infectivity

To determine whether, in the context of EBOV infection, VP24 can traffic to the nucleus, we transfected Huh7 cells with plasmids that express Flag-tagged IMPA3 or IMPA7 and then challenged with EBOV. We found both IMPA3 and IMPA7 were present almost exclusively in the cell nucleus. However, strong nuclear VP24 expression was only evident in IMPA7-transfected cells, demonstrating the capacity of VP24 to enter the nucleus in infected cells and suggesting that specific IMPA7 association with VP24 plays a significant role in its nuclear localization (Fig. 1).

**Figure 1:**
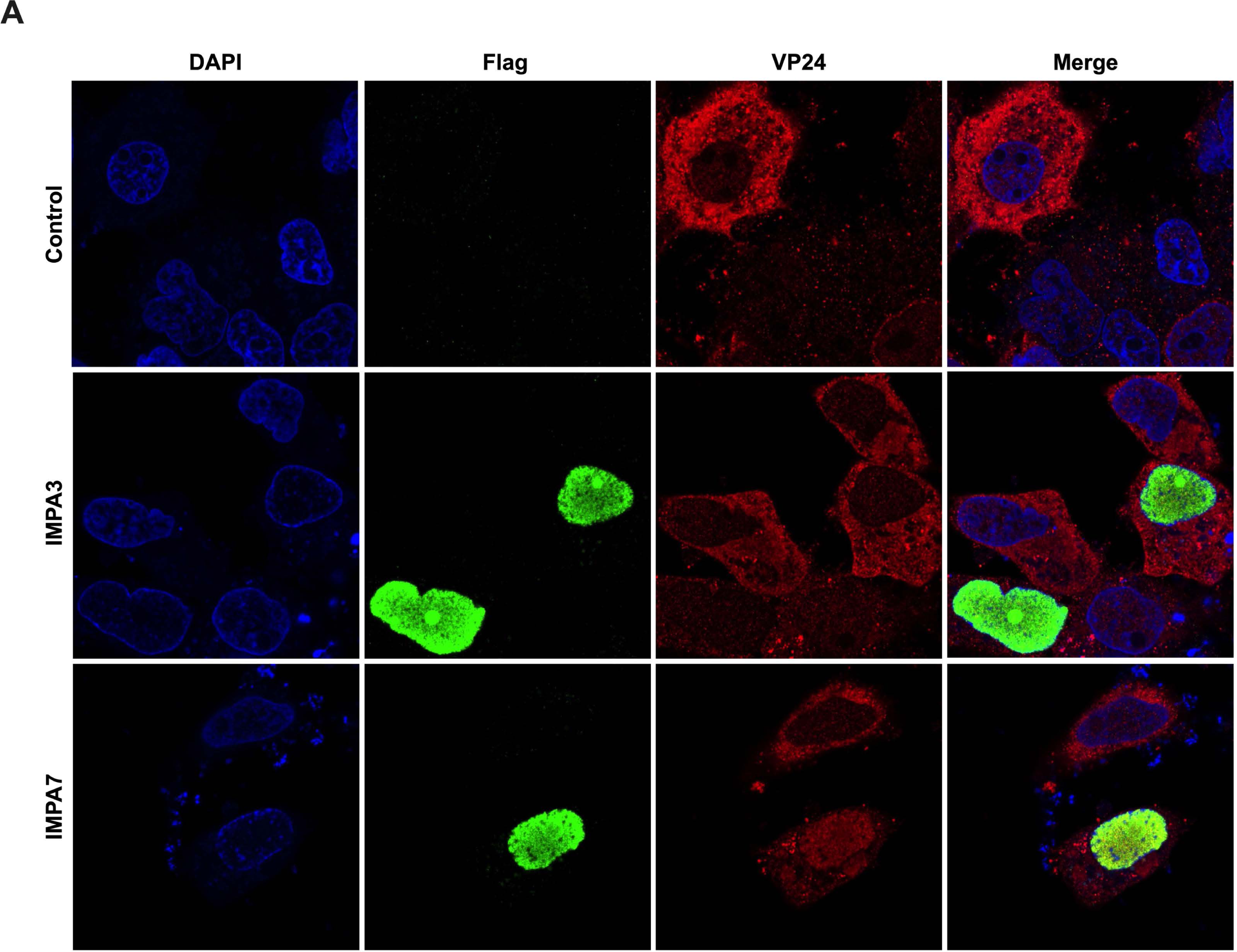
VP24 Localizes with Nuclear IMPA7 in EBOV Infection. Representative confocal images of VP24 in EBOV-infected and IMPA-transfected cells are provided. Huh7 cells transfected with Lipofectamine 3000 and plasmids for Flag-tagged IMPA3 or IMPA7. Control indicates Huh7 cells treated with lipofectamine 3000 alone. Cells were stained with anti-flag (green) or VP24 protein (red) and cell nuclei with Hoeschst 33342. All images are single plane confocal images through the middle of the cell body, and all channel intensities were normalized to the control using ImageJ.

To examine the significance of VP24 nuclear trafficking during EBOV infection, we utilized an EBOV transcription and replication competent virus-like particle system (trVLP) [20, 42]. This life cycle modeling system allows for the examination of crucial steps in the EBOV life cycle without the need for a high-containment laboratory (Fig 2A). A key component of this system is a tetracistronic minigenome that encodes for *Renilla* luciferase as well as the EBOV proteins VP40, GP, and VP24. The minigenome also includes the EBOV leader and trailer sequences, three of the EBOV non-coding regions (NCRs) and a T7 promoter. At passage 0 (P0), trVLPs are generated in producer cells by transfecting Huh7 cells with the tetracistronic minigenome, T7 RNA polymerase, and the EBOV polymerase complex proteins VP30, VP35, NP and L. The T7 RNA polymerase is required for initial mRNA transcription from the minigenome, while the EBOV polymerase complex is responsible for replication of the minigenome RNA. Measured by reporter assay, *Renilla* luciferase activity at P0 serves as a readout for viral genome replication and gene expression. As a control for reporter activity, we also include a condition where L, the catalytic component of polymerase complex, is replaced with empty vector (-L). Following passage of trVLPs onto target cells expressing the EBOV polymerase complex at P1, *Renilla* luciferase activity can be analyzed to measure infectivity of trVLPs generated in P0.

**Figure 2:**
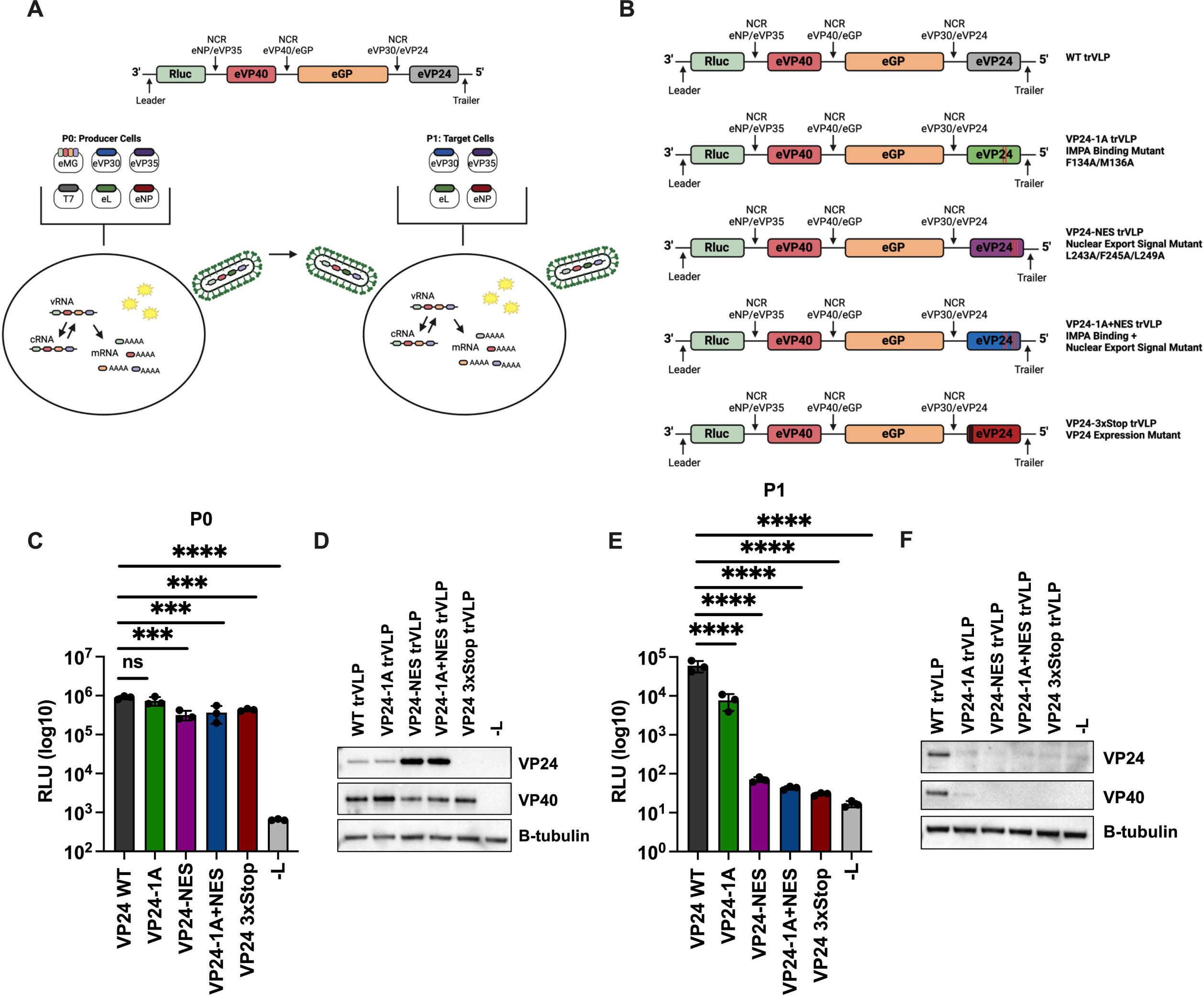
The VP24-IMPA Interface and VP24 NES are Important for Transcription and Replication Competent Virus-Like Particle (trVLP) Infectivity. A) Schematic summary of the tetracistronic minigenome reporter and trVLP assay. The tetracistronic minigenome encodes for *Renilla* luciferase and viral proteins VP40, GP, and VP24. The genes are separated by non-protein coding regions (NCR) from the indicated EBOV genes. The minigenome plasmid also includes the EBOV leader and trailer sequences, and the T7 RNA polymerase promoter. The P0 transfection includes the minigenome plasmid (eMG) and plasmids that express T7 RNA polymerase (T7), EBOV VP30 (eVP30), EBOV VP35 (eVP35), EBOV L (eL) an EBOV NP (eNP). Reconstitution of the viral polymerase complex after transfection leads to the replication of the trVLP genome, resulting in production of negative-sense genomic vRNA and positive sense antigenome RNA (cRNA) and in production of mRNAs. Minigenome reporter activity is analyzed by luciferase assay to assess viral genome replication and gene expression. P0 supernatants containing trVLPs are harvested and passaged onto target cells that have been transfected with the EBOV polymerase complex plasmids (P1). Examination of minigenome reporter activity at P1 provides a measure of trVLP infectivity. B) Schematic representation of the different wildtype (WT) and mutant VP24 tetracistronic minigenomes. The mutants are as follows: VP24 cluster 1A IMPA binding mutant (VP24-1A), VP24 nuclear export signal mutant (VP24-NES), VP24 IMPA binding mutant + NES mutant (VP24-1A+NES), and VP24 expression mutant (VP24-3xStop). C) Analysis of P0 reporter activity. A P0 transfection was performed in Huh7 cells with the indicated minigenomes. As a control, empty vector was transfected in place of L (-L). Seventy-two hours post-transfection, trVLP supernatants were collected and passaged onto target cells. The remaining P0 cells were lysed to measure *Renilla* luciferase activity, represented as relative light units (RLU). E) Analysis of P1 reporter activity. P1 transfected target cells were infected with the indicated P0 trVLP supernatants. Seventy-two hours post trVLP infection, P1 cells were lysed to measure *Renilla* luciferase activity. D, F) Western blots of reporter assay lysates. Lysates from triplicate P0 (D) or P1 (F) samples were pooled and analyzed by western blot for VP24 and VP40 expression. Expression of β-tubulin is shown as a loading control. A, B) Figure was created with BioRender.com. C, E) Error bars represent the standard error for triplicate samples. **** denotes p-value ≤ 0.001. *** denotes p-value ≤ 0.001. NS denotes p-value > 0.05.

We generated tetracistronic minigenomes with mutations in VP24 targeting the previously described cluster 1A residues of the IMPA interface (VP24-1A) and the NES (VP24-NES) (Fig 2B). VP24-1A has the mutations F134A/M136A [24] and VP24-NES contains the mutations L243A, F245A, L249A that were previously described to disable VP24 nuclear export [38]. We also generated a combined KPNA binding plus NES mutant (VP24-1A+NES) [38]. As an additional control, we incorporated three stop codons after the VP24 start codon to abolish VP24 expression while maintaining minigenome length (VP24-3xStop), as has been reported for prior VP24 studies [20].

When tested in the trVLP assay, the wildtype and mutant minigenomes, including VP24-3xStop, exhibited similar P0 reporter activities, with slight reductions for both NES-mutants and VP24-3xStop (Fig 2C). Western blots of lysates from P0 demonstrated expression of the wildtype and mutant VP24s (Fig 2D). Interestingly, VP24-NES and VP24-1A+NES expressed elevated levels of VP24 compared to VP24 WT. As expected, VP24-3xStop did not express VP24. The minigenome-encoded viral protein VP40 was also readily detected in each sample, with modest reductions seen once again for both NES-mutants and VP24-3xStop, supporting the results of the reporter assay (Fig. 2D). These data indicate that VP24 and the VP24 trafficking functions have at most modest impacts on viral RNA synthesis and gene expression at P0.

Supernatants containing trVLPs generated from the above P0 transfection were harvested and passed onto target cells to assess the impact of the nuclear trafficking mutations on trVLP infectivity. At P1, compared to VP24 WT, VP24-1A exhibited a roughly tenfold reduction in reporter activity (Fig 2E). In contrast, reporter activity was reduced by approximately 1000-fold for VP24-NES and VP24-1A+NES. This reduction was similar to that observed for the VP24-3xStop mutant and the -L control. While VP24 and VP40 expression could be detected in the VP24 WT P1 lysates, VP24 could not be detected for any of the mutants and only a faint band for VP40 was detected for the VP24-1A mutant (Fig 2F). At P1, the VP24-1A, VP24-NES, and VP24-1A+NES trVLP infected cells also exhibited greatly reduced levels of VP40 mRNA and the EBOV 5’ trailer, as measured by reverse transcription-quantitative PCR (RT-qPCR), demonstrating a reduction in viral genome transcription and replication for these mutants (S1A-B Fig). Furthermore, levels of viral proteins present in concentrated P0-derived trVLPs were reduced in VP24-1A, VP24-NES, and VP24-1A+NES trVLPs (S1C Fig). Together, these results suggest that the VP24 nuclear trafficking signals play an important role in trVLP infectivity and production of infectious viral particles, with the VP24-NES mutants showing the strongest negative impact on trVLP infectivity.

### VP24 Nuclear Export is not Required for Inhibition of STAT1-dependent Gene Expression or IFN-β Promoter Driven Gene Expression

We next sought to determine how the mutations affect inhibition of IFN responses. Co-immunoprecipitation experiments confirmed that both VP24 WT and VP24-NES interact with IMPA6 (Fig 3A). As anticipated, the IMPA binding mutants VP24-1A and VP24-1A+NES lost interaction with IMPA6. As in prior studies, inhibition correlated with IMPA interaction. VP24 WT and VP24-NES significantly suppressed inhibition of IFN-stimulated response element (ISRE) promoter drive firefly luciferase expression, while VP24-1A and VP24-1A+NES mutants were less inhibitory (Fig 3B). In contrast, VP24 WT and the VP24 mutants each suppressed Sendai virus (SeV)-induced activation of an IFN-β promoter driven firefly luciferase reporter (Fig 3C). These data correlate VP24-IMPA interaction with inhibition of IFN signaling, demonstrate that VP24 nuclear trafficking signals are not required for VP24-mediated suppression of IFN-β promoter activity and indicate that the infectivity defect of the VP24-NES and VP24-1A+NES mutants are independent of VP24 IFN-inhibitory functions.

**Figure 3:**
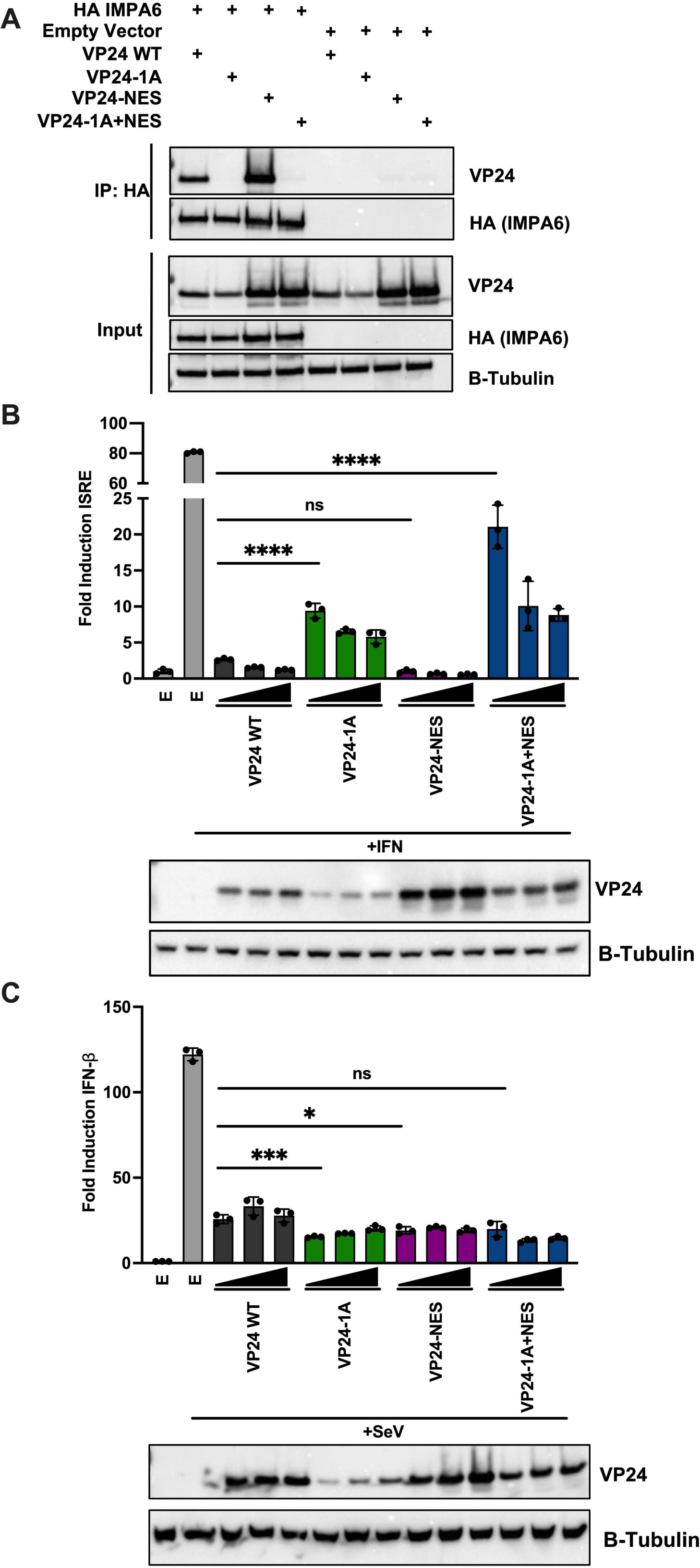
VP24-NES Inhibits STAT1-dependent Gene Expression and IFN-β Promoter Activity. A) A co-immunoprecipitation assay was performed in 293T cells to assess interaction between HA-tagged IMPA6 and WT or mutant VP24s. HA-IMPA6 was immunoprecipitated and probed for interaction with VP24 by western blot. Empty pCAGGS vector was used as a control for non-specific interaction. Expression of β-tubulin is shown as a loading control for the input samples. B) Analysis of IFN-stimulated response element (ISRE) promoter-firefly luciferase reporter activity after treatment with universal type I IFN. 293T cells were transfected with the ISRE luciferase reporter and VP24 WT or mutant constructs. Empty vector pCAGGS (E) served as a control. Reporter activity is represented as fold induction of reporter relative to empty vector, mock-treated cells. C) Analysis of IFN-β-firefly luciferase reporter activity after Sendai virus (SeV) infection. 293Ts were transfected with the IFN-β luciferase reporter and VP24 WT or mutant constructs. Empty vector pCAGGS (E) was used as a control. Reporter activity is represented as fold induction of reporter relative to empty vector mock. B-C) Reporter assay lysates were analyzed by western blot for VP24 expression. Expression of β-tubulin is shown as a loading control. Error bars represent the standard deviation for triplicate samples. **** denotes p-value ≤ 0.001. *** denotes p-value ≤ 0.001. * ≤ denotes p-value 0.05. ns denotes p-value > 0.05.

### VP24 Nuclear Import and Export Signals are Functional

To directly assess the function of VP24 nuclear trafficking signals, the subcellular distribution of VP24 was examined in the context of the trVLP assay. Following a P0 transfection, cytoplasmic and nuclear fractions were prepared and examined by western blot. HDAC2 and β-tubulin served as nuclear and cytoplasmic markers. VP24 WT and VP24-NES were detected in both the cytoplasmic and nuclear fractions, beginning at 72 hours post-transfection for VP24 WT and 48 hours post-transfection for VP24-NES (Fig 4A). In contrast, VP24-1A and VP24-1A+NES were predominantly cytoplasmic, with a slight signal in the nuclear fraction (Fig 4A). Immunofluorescence microscopy was then used to calculate the average nuclear fluorescence over the average cytoplasmic fluorescence (fn/c) of VP24 after a P0 transfection. While VP24 WT exhibits some nuclear localization, it predominantly localizes in the cytoplasm, where it forms distinct perinuclear puncta (Fig 4B, C). Compared to VP24 WT, VP24-1A localization is significantly less nuclear, demonstrating that disrupting the VP24-IMPA interaction hinders VP24 nuclear import. In contrast, VP24-NES is significantly more nuclear than VP24 WT, suggesting impaired nuclear export. Despite its pronounced nuclear localization, VP24-NES is still detected in the cytoplasm where it exhibits a more diffuse distribution than VP24-WT or VP24-1A. VP24-1A+NES is more nuclear than VP24-1A but more cytoplasmic than the VP24-NES mutant, resulting in an fn/c value similar to VP24 WT. Previous *in vitro* binding studies have demonstrated that VP24-1A retains weak binding to IMPA [25]. Therefore, it is possible that VP24-1A inefficiently traffics to the nucleus and the addition of the NES mutations leads to some accumulation of VP24-1A+NES in the nucleus.

**Figure 4:**
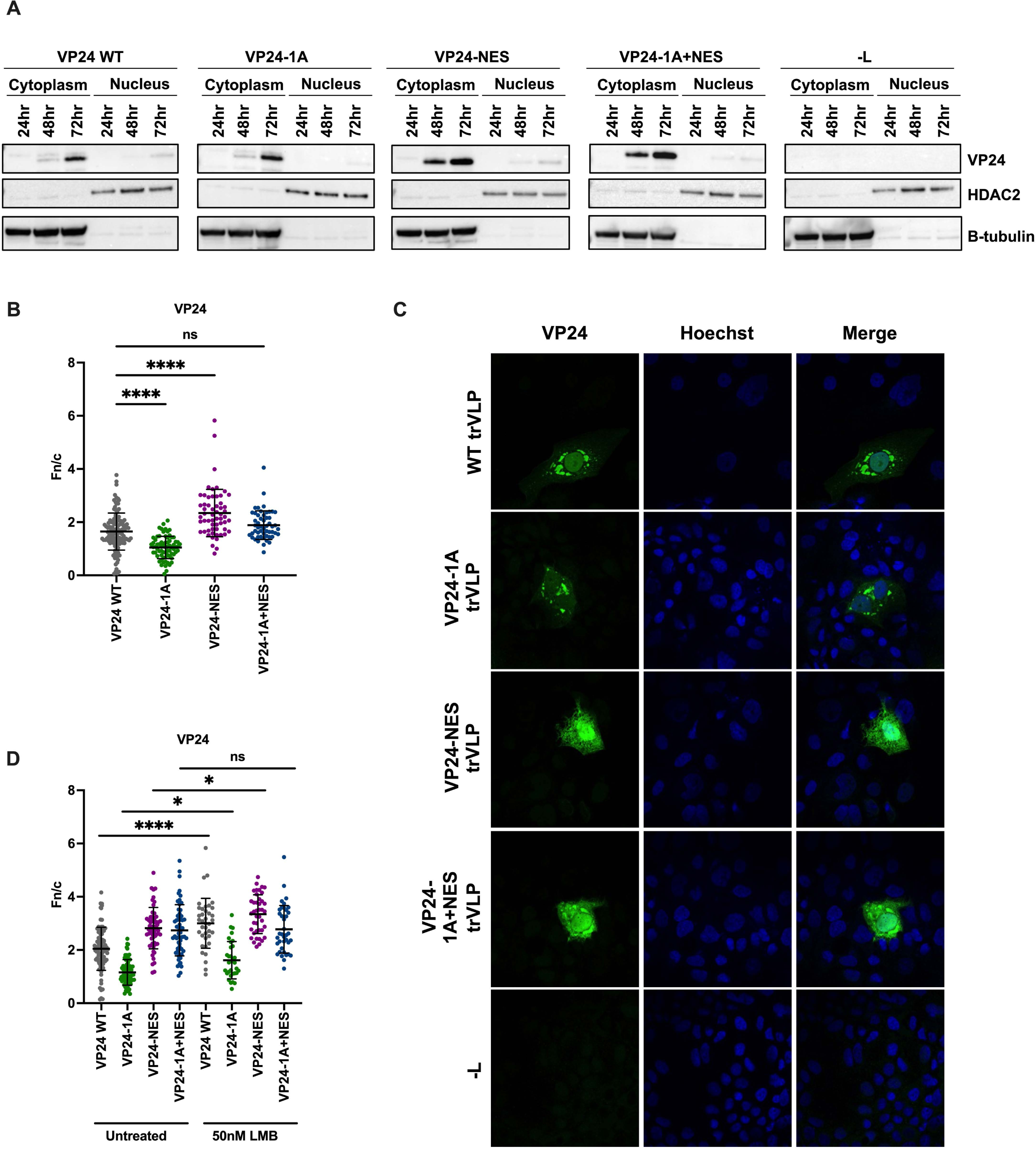
VP24 Possesses Functional Nuclear Trafficking Signals. A) Western blot analysis of VP24 levels in nuclear and cytoplasmic fractions at P0. A P0 transfection was performed in Huh7 cells with VP24 WT or mutant minigenomes. –L indicates a P0 transfection with the VP24 WT minigenome in which the L expression plasmid was omitted. Cells were harvested at the indicated time points post-transfection, and nuclear and cytoplasmic fractions were prepared. VP24 expression was assessed by western blot. HDAC2 expression served as a control for nuclear fractions and β-tubulin expression served as a control for cytoplasmic fractions. B-C) Analysis of VP24 localization by immunofluorescence microscopy. Seventy-two hours after a P0 transfection with the indicated minigenomes, Huh7 cells were fixed and stained for VP24 and nuclei. B) Average nuclear fluorescence over average cytoplasmic fluorescence (fn/c) for VP24. C) Representative confocal images of VP24 localization. D) Quantification of VP24 fn/c following treatment of transfected cells with 50nM leptomycin B (LMB). B, D) Data are represented as mean fn/c for each minigenome ± SD. **** denotes p-value ≤ 0.001. * ≤ denotes p-value 0.05. ns denotes p-value > 0.05.

To further examine the functionality of the VP24 NES, we examined VP24 localization after treatment with leptomycin B (LMB), an inhibitor of the nuclear export protein CRM1, after P0 transfection. LMB increased VP24 WT nuclear localization as compared to the untreated VP24 WT control (Fig 4D). In contrast, treatment with LMB had only a modest impact on the VP24 mutants. Together, these results establish that in the context of the trVLP assay VP24 has functional nuclear trafficking signals.

### The VP24 Nuclear Export Signal is Important for EBOV Nucleocapsid Formation

The impact of VP24-WT versus mutant overexpression on tetracistronic minigenome reporter activity was examined. Overexpression of VP24 WT resulted in a greater than tenfold inhibition of reporter activity when compared to empty vector control (Fig 5A). VP24-1A also inhibited reporter activity, yielding an almost 100-fold reduction. In contrast, VP24-NES and VP24-1A+NES caused only a modest reduction in reporter activity with luciferase levels remaining at least 10-fold higher than the VP24-WT condition, suggesting that the VP24-NES mutants are defective in nucleocapsid condensation. Western blotting of lysates from the P0 cells demonstrated comparable expression of each of the over-expressed VP24s and levels of VP40 expression from the trVLP genome paralleled the reporter gene data (Fig 5B).

**Figure 5:**
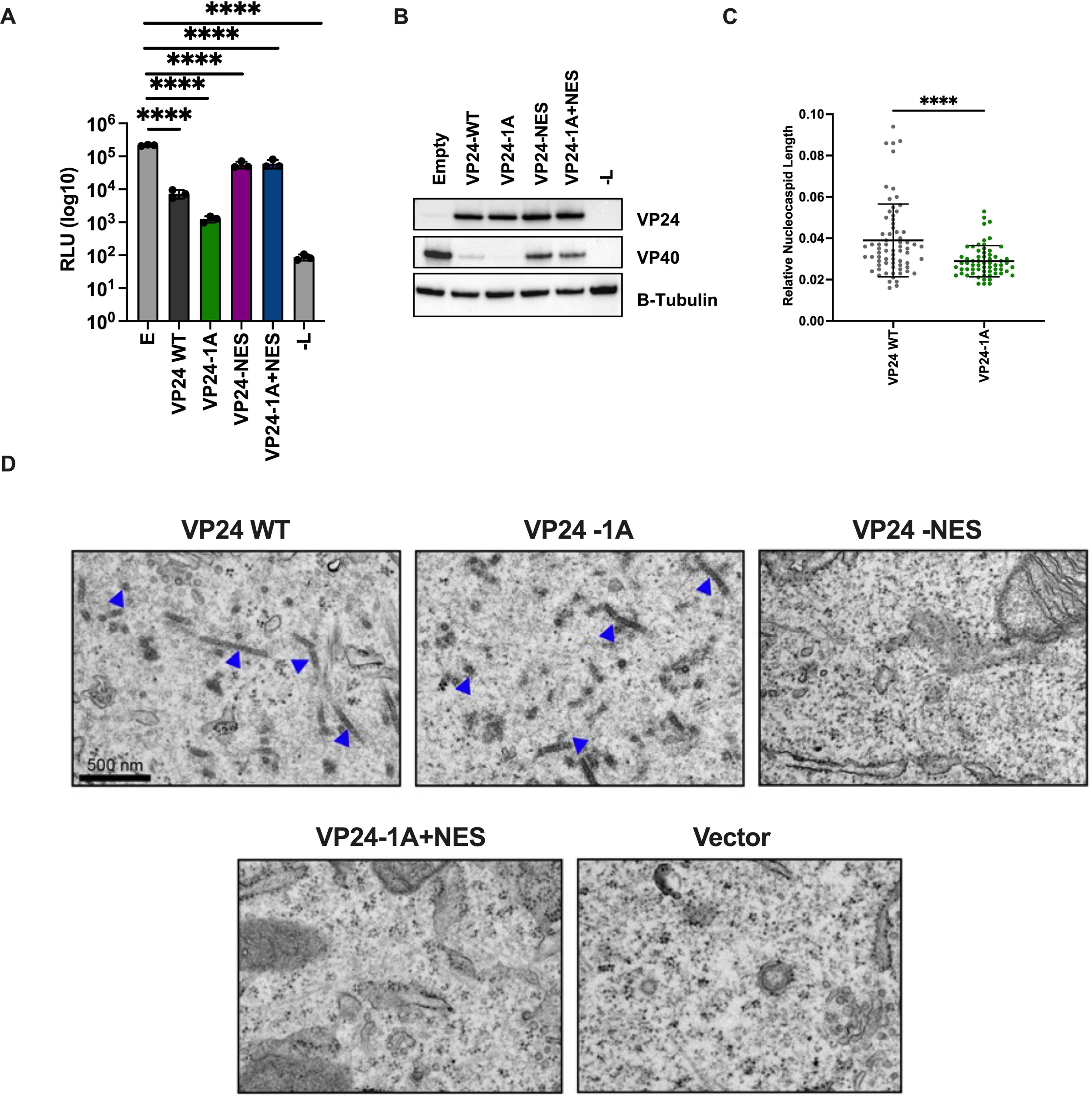
VP24 NES is Required for Nucleocapsid-like Structure Assembly. A) Analysis of VP24-mediated inhibition of P0 minigenome reporter activity. In addition to a typical P0 transfection in Huh7 cells, VP24 WT or mutant proteins were overexpressed. Empty vector (E) was included as a control. Seventy-two hours post-transfection cells were lysed to measure *Renilla* luciferase activity. Error bars represent standard error for triplicate samples. B) Western blot analysis of minigenome reporter assay lysates. Triplicate wells from P0 reporter assay lysates were pooled and analyzed by western blot for VP24 and VP40 levels. Expression of β-tubulin is shown as a loading control. C) Quantification of relative nucleocapsid length. Huh7 cells were transfected with NP, VP35, and the indicated VP24 plasmids. Empty pCAGGS vector in place of VP24 was included as a control. Twenty-four hours post-transfection, cells were fixed and processed for electron microscopy analysis. The FIJI image processing program was used to calculate the relative nucleocapsid length for samples exhibiting electron-dense nucleocapsid-like structures. Error bars represent the standard deviation of more than 50 nucleocapsid-like structures analyzed from the VP24 WT and VP24-1A image sets. **** denotes p-value ≤ 0.001. D) Representative electron microscopy images assessing the presence of nucleocapsid-like structures. Arrows point to nucleocapsid-like structures. Scale bar = 500 nm, valid for all EMD frames.

VP24 co-localizes with NP and VP35 in inclusion bodies that form during viral infection [17, 35]. In the context of a P0 transfection, VP24 WT and VP35 colocalized in perinuclear puncta, suggesting these puncta are viral inclusion bodies. VP24-NES and VP24-1A+NES partially colocalized with VP35, however, a substantial amount of each mutant remained outside of the inclusions in both the nucleus and the cytoplasm (S2 A-B Fig). The altered cytoplasmic distribution of VP24-NES and VP24-1A+NES could reflect altered interaction with NP and VP35, although it might also reflect the higher expression seen with these mutants.

Overexpression of the essential nucleocapsid components NP, VP35, and VP24 is sufficient to form nucleocapsid-like structures that, when examined by electron microscopy, closely resemble the nucleocapsids formed during EBOV infection [11, 17]. When VP24 WT or VP24-1A were co-expressed with NP and VP35, formation of nucleocapsids was observed (Fig 5C-D), suggesting that the VP24-IMPA interaction is not required for the formation of these structures. However, nucleocapsid-like structures formed in the presence of VP24-1A were slightly shorter on average than those observed with VP24 WT (Fig 5C). This suggests that the VP24-IMPA interaction is not required for nucleocapsid formation, but the cluster 1A IMPA binding residues or VP24-IMPA interaction contribute to this process. In contrast to VP24 WT and VP24-1A, nucleocapsid-like structures were not detected in cells co-expressing VP24-NES or VP24-1A+NES with NP and VP35 (Fig. 5D). This suggests a critical role for VP24 nuclear export or for the residues within the NES in nucleocapsid-like structure assembly.

### VP24 Nuclear Trafficking is not Sufficient for the Production of Infectious Particles

To address whether loss of NES activity explains the viral assembly/infectivity defects in the NES mutants, we generated new VP24 mutant minigenomes (Fig S3). In one, we exchanged the VP24 NES with the NES from the protein kinase A inhibitor (PKI) protein (VP24-PKI NES). We also generated a minigenome in which we mutated the residues surrounding the VP24 NES to alanine (E244A/N246A/S247A/S248A), keeping the key NES residues intact (VP24-ENSS). To further assess the impact of VP24 nuclear import on trVLP infectivity, we mutated the cluster 3 residues (L201A/E203A/P204A/D205A/S207A) of VP24 that also contribute to IMPA interaction (VP24-3) [24].

To determine the impact of the mutations, VP24 localization was assessed through immunofluorescence microscopy. VP24-3, like VP24-1A, exhibited decreased nuclear localization as reflected in a lower fn/c value when compared to VP24 WT (Fig 6A). Compared to VP24-NES, both the VP24-ENSS and VP24-PKI NES exhibited reduced nuclear localization with fn/c values comparable to VP24 WT, demonstrating functional VP24 nuclear export for these mutants.

**Figure 6:**
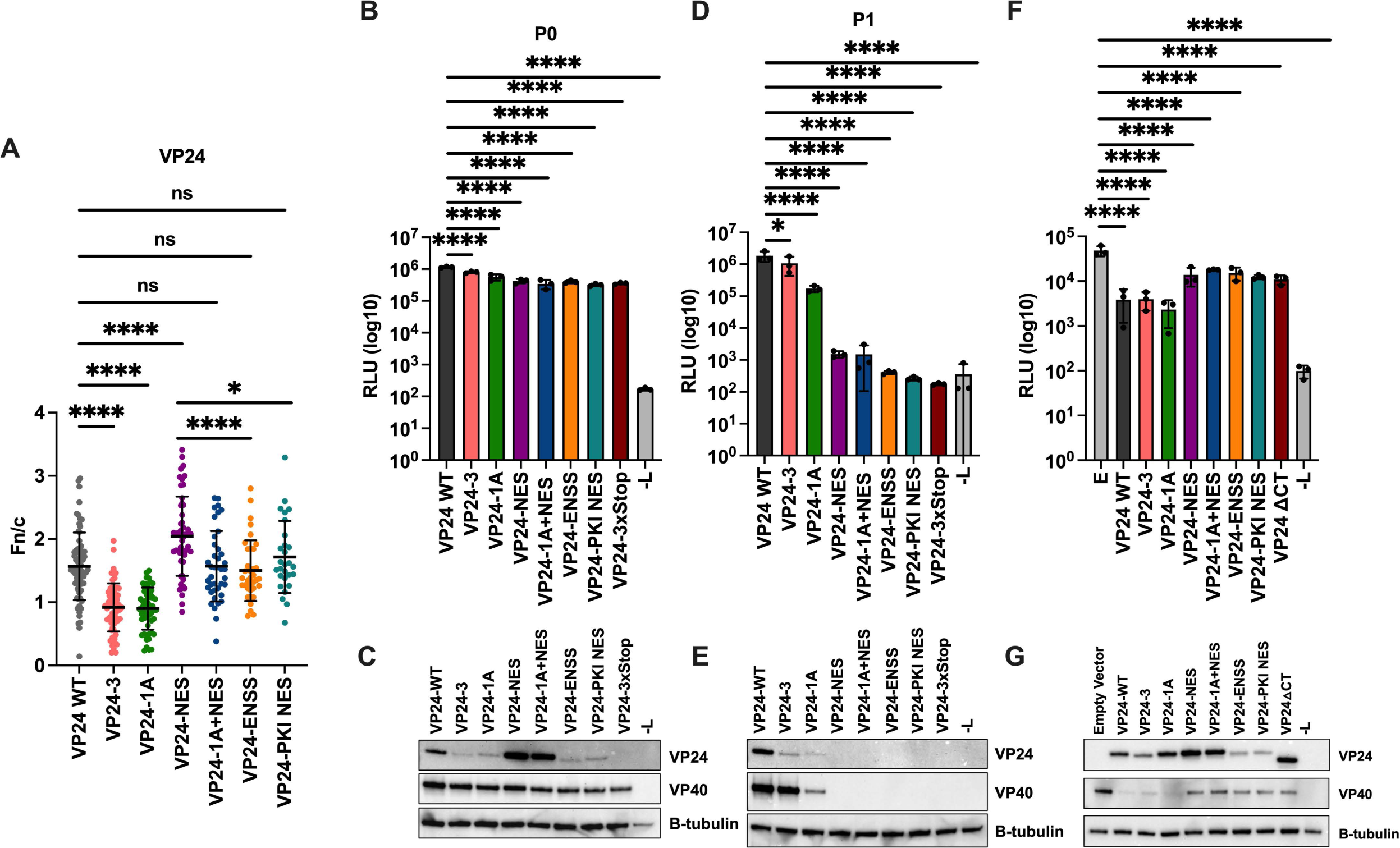
Nuclear Trafficking of VP24 is Not Sufficient for the Production of Infectious trVLPs. A) Quantification of VP24 fn/c at P0. A P0 transfection was performed in Huh7 cells with the indicated minigenomes. In addition to VP24 WT, VP24-1A, VP24-NES and VP241A+NES; mutant minigenomes included in this assay are a VP24 cluster 3 IMPA binding mutant (VP24-3), a VP24 mutant with the Protein Kinase A Inhibitor (PKI) NES instead of the VP24 NES (VP24-PKI NES), a VP24 mutant where the residues surrounding the NES were mutated to alanine (L201A/E203A/P204A/D205A/S207A) (VP24-ENSS), and a VP24 mutant where residues 232-252 were deleted (VP24ΔCT). Seventy-two hours post-transfection cells were fixed and stained as described in Figure 3. Data are represented as mean fn/c for each minigenome ± SD. B) Analysis of P0 reporter activity. A P0 transfection was performed in Huh7 cells with the indicated minigenomes. –L indicates a P0 transfection with the VP24 WT minigenome in which the L expression plasmid was omitted. trVLP supernatants were collected 72 hours post-transfection and used to infect target cells. The remaining cells were lysed to measure *Renilla* luciferase activity. D) Analysis of P1 minigenome reporter activity. P0 trVLP supernatants were used to infect P1 transfected target cells. Seventy-two hours post trVLP infection, cells were lysed to measure *Renilla* luciferase activity. C, E) Lysates from P0 (C) or P1 (E) minigenome reporter assays were pooled and analyzed by western blot for VP24 and VP40 expression. Expression of β-tubulin is shown as a loading control. F) Analysis of VP24-mediated inhibition of minigenome reporter activity. The indicated VP24 plasmids were overexpressed during a typical P0 transfection. Empty vector (E) and a – L transfection were included as controls. Seventy-two hours post-transfection cells were lysed to measured *Renilla* luciferase activity. G) Western blot analysis of VP24 overexpression reporter assay lysates. Assay lysates from triplicate wells were pooled and analyzed by western blot for VP24 and VP40 expression. Expression of β-tubulin is shown as a loading control. Error bars represent the standard error for triplicate samples. * *** denotes p-value ≤ 0.001. * ≤ denotes p-value 0.05. NS denotes p-value > 0.05.

At P0 in the trVLP assay, all the mutant minigenomes exhibited reporter activity similar to VP24 WT (Fig 6B). Western blots of the reporter assay lysates demonstrated that VP24-3, VP24-ENSS, and VP24-PKI NES expressed to similar levels as VP24-1A but to somewhat lower levels than VP24 WT (Fig 6C). As with the reporter assay, VP40 expression was similar for all mutant minigenomes, further demonstrating that viral gene expression is not impacted by these mutations. At P1, the VP24-3 mutant exhibited a very modest reduction trVLP infectivity, compared to the 10-fold reduction for the VP24-1A mutant (Fig 6D). This was also reflected in the western blots of the P1 lysates for VP24-3 and VP24-1A (Fig 6E). P1 infectivity was reduced by almost 1000-fold for VP24-ENSS and VP24-PKI NES (Fig 6D), despite these mutants exhibiting similar localization as VP24 WT (Fig 6A). Similarly, VP24 and VP40 expression were not detectable in the VP24-ENSS and VP24-PKI NES P1 lysates (Fig 6E). This reduction in infectivity is similar to that observed with VP24-NES, suggesting that VP24 nuclear export is not sufficient for the production of infectious particles.

Finally, we examined VP24-mediated inhibition of minigenome reporter activity as a proxy for condensation of nucleocapsid structures. VP24 WT, VP24-3, and VP24-1A inhibited minigenome reporter activity by more than 10-fold when overexpressed at P0, suggesting that VP24 nuclear import is not required for nucleocapsid condensation (Fig 6F). In contrast, overexpressed VP24-ENSS and VP24-PKI NES did not inhibit minigenome reporter activity to the same degree as VP24 WT and closely resembled VP24-NES, suggesting that nuclear trafficking is not sufficient to promote nucleocapsid condensation. Therefore, the VP24 C-terminus contributes by other mechanisms to nucleocapsid condensation. Given the key role for the C-terminus in inhibition of transcription and replication, we tested an additional VP24 mutant, in which the crucial C-terminus of VP24 was removed (VP24-ΔCT). Similar to VP24-NES, VP24-1A+NES, VP24-ENSS, and VP24-PKI NES, VP24-ΔCT did not inhibit reporter activity to the same degree as VP24 WT or the IMPA binding mutants. In these assays, overexpressed VP24-ENSS and VP24-PKI NES expressed to slightly lower levels than VP24 WT in the lysates of the P0 cells, while VP40 expression reflected the reporter gene data (Fig 6G). Together, these results suggest that the VP24 NES residues are important for nucleocapsid assembly; however, VP24 nuclear export is not sufficient for nucleocapsid condensation.

## Discussion

EBOV VP24 is a multifunctional protein with important roles in viral nucleocapsid assembly and innate immune antagonism. In this study, we explored the importance of VP24 in the EBOV life cycle by focusing on the VP24-IMPA interface and the VP24 NES. As an innate immune antagonist, VP24 prevents STAT1-IMPA interaction by binding to armadillo repeats (ARMS) 8-10 of IMPA5, IMPA6 and IMPA7, a region outside of the classical NLS binding used by most IMPA cargo [23, 24]. The VP24-IMPA interface is critical for VP24-mediated inhibition of STAT1-dependent gene expression. Consistent with its functional significance, VP24-IMPA interaction is highly conserved among the *Orthoebolavirus* species and in LLOV, though the strength of this interaction can differ between VP24s from different viruses [25, 26, 31]. VP24 has a pyramidal structure and in the IMPA-VP24 crystal structure, three clusters of amino acid residues in VP24 made contact with IMPA [24, 43]. Cluster 1 and 3 amino acid residues are highly conserved among members of the *Orthoebolavirus* genus. Furthermore, mutations in these clusters, such as the cluster 1A mutant (F134A, M136A) and the cluster 3 mutant (L201A, E320A, P204A, D205A, S207A) lead to impaired VP24-IMPA interaction and inhibition of IFN signaling [24].

While well characterized for its role in inhibiting IFN signaling, the interaction between VP24 and IMPA also suggests that VP24 nuclear import is possible. However, neither VP24-IMPA interaction nor VP24 nuclear trafficking had been demonstrated in EBOV infected cells. Rather, VP24 has a cytoplasmic steady-state localization during infection [8, 35]. In transfected cells, over-expression of NPI-1 subfamily member IMPA6 can relocate an otherwise cytoplasmic GFP-VP24 protein into the nucleus, suggesting VP24 can traffic to the nucleus via IMPA [38, 44]. Here, we demonstrated that expression of another NPI-1 subfamily member, IMPA7, dramatically increased VP24 nuclear localization during EBOV infection (Fig 1). In contrast, expression of IMPA3 did not promote VP24 nuclear localization in infected cells. This provides evidence of functional IMPA7-VP24 interaction during infection and demonstrates that VP24 can traffic to the nucleus via this interaction. In the context of the trVLP assay, we determined that mutations in cluster 1A or 3 that hinder VP24-IMPA interaction lead to impaired nuclear import of VP24 (Fig 4 and Fig 6A). We also demonstrated that the VP24 NES is functional in the context of the trVLP system, observing increased nuclear accumulation of VP24 following mutation of the VP24 NES or treatment with LMB (Fig 4B-D). This is consistent with the prior report that showed that mutating the NES redirects VP24 to the nucleus [38]. Cumulatively, these data establish that VP24 traffics to the nucleus via IMPA interaction and has a functional NES.

Prior studies demonstrated that transfected VP24 can also suppress IFN-β promoter activity, but it was unclear whether this inhibition relies upon the VP24-IMPA interaction [22]. We demonstrate that VP24-IMPA interaction was not required for the suppression of SeV induced IFN-β promoter driven reporter activity in our assays (Fig 3B-C). This suggests that an alternative mechanism may be employed by VP24 to suppress the IFN-β promoter. In contrast to our findings, a recent paper demonstrated that VP24-IMPA interaction is required to suppress IFN-β promoter driven reporter activity induced by MDA5 or constitutively active RIG-I expression [33]. This difference in suppression could be attributed to the different VP24-IMPA interaction mutants used for these experiments (VP24-1A vs VP24-3). Additional studies are required to determine the exact mechanism of VP24-mediated suppression of the IFN-β promoter.

In addition to its role as an innate immune antagonist, VP24 is critical for nucleocapsid assembly and the production of infectious viral particles [11, 19, 21]. Previous studies have shown that knockdown of VP24 during EBOV infection leads to the accumulation of malformed nucleocapsids and impaired viral release [21]. Studies where EBOV has been adapted to mouse or guinea pigs provide evidence for a role of VP24 in nucleocapsids formation. Mutations in VP24 are associated with the acquisition of virulence in mouse- and guinea pig-adapted strains of EBOV [45–47]. In the case of mouse-adapted EBOV, mutations in both NP and VP24 were associated with increased virulence in mice [45]. However, VP24 adaptive mutations alone had little impact on viral replication in IFN treated RAW 264.7 cells, making it unclear whether these adaptive mutations impact STAT1 nuclear trafficking and leaving open the possibility that other VP24 functions are affected [45]. In infected primary guinea pig macrophages, wildtype EBOV did not produce viral nucleocapsids whereas a recombinant EBOV possessing guinea pig adaptive mutations in VP24 generated normal nucleocapsids [47]. Also supporting the role of VP24 in nucleocapsid formation and infectivity, a VP24 mutant with impaired NP interaction exhibited reduced trVLP production, likely due to impaired nucleocapsid assembly [48]. Using a monocistronic minigenome system in which all the EBOV proteins are overexpressed, it was shown that mutations in a VP24 YXXL motif led to shorter nucleocapsid-like structures, impaired nucleocapsid transport, and reduced trVLP infectivity [49]. Additionally, the authors were unable to rescue recombinant YXXL mutant EBOV, suggesting that these mutations severely impact virus production [49].

Given the importance of VP24 for EBOV infectivity, we examined how mutations in the VP24-IMPA interface or NES impacted nucleocapsid assembly and trVLP infectivity. We observed that disrupting the VP24-IMPA interaction led to a modest reduction in infectivity at P1 and shortened nucleocapsid-like structures (Fig 2E, Fig 5D, and Fig 6D). While cluster 1A and 3 mutations equally impeded VP24 nuclear import, we observed lower infectivity for VP24-1A as compared to VP24-3 (Fig 6D). This suggests that the cluster 1A mutations may impact other VP24 functions or interactions. Nevertheless, the reduced infectivity for both mutants further serves to underscore the importance and complexity of the VP24-IMPA interface in the EBOV life cycle.

Our studies also revealed that mutating VP24 NES residues had a profound effect on the production of trVLP particles, leading to severely impaired trVLP infectivity (Fig 2E). Upon closer examination, we discovered that mutating the NES impairs the ability of VP24 to inhibit minigenome reporter activity, suggesting that the NES mutant has impaired capacity to interact with the viral RNA synthesis machinery. We also discovered a loss of nucleocapsid formation for the NES mutant when expressed with NP and VP35 (Fig 5A-C). Together, this establishes that the defect in infectivity for VP24-NES trVLPs is due to impaired nucleocapsid assembly, highlighting a previously unknown function for the VP24 NES residues.

VP24 N- or C-terminal truncations as small as 5 amino acid residues can also inhibit nucleocapsid formation, demonstrating that small changes to the termini of VP24 can drastically alter function [18]. In the case of the 5 amino acid C-terminal deletion, one of the NES residues mutated in our study was removed [18]. Therefore, it is unclear whether the phenotype of this mutant was due to loss of NES function or for other reasons. Even the addition of a Flag-tag to the N- or C-terminus of VP24 can impair nucleocapsid assembly [13, 17]. As a result, all of our assays were performed using untagged VP24. This also precluded us from adding an NLS or NES sequence to VP24. With this is mind, we generated new VP24 mutants to determine the importance of VP24 nuclear trafficking for trVLP infectivity (S3 Fig). To examine the importance of the NES region, we kept the NES residues intact but mutated the residues surrounding the NES to alanine (E244A/N246A/S247A/S248A) (VP24-ENSS). We also generated a mutant where we replaced the VP24 NES with the NES from the Protein Kinase A Inhibitor protein (PKI-NES). The PKI NES and VP24 NES are both leucine-rich NESs that are very similar in size and were used to minimize alterations to the VP24 C-terminus. Despite exhibiting functional nuclear trafficking, VP24-ENSS and VP24-PKI NES trVLP infectivity was impaired (Fig 6A-E). These mutants were also unable to inhibit minigenome reporter activity when overexpressed (Fig 6F), suggesting the impaired infectivity is due to defective nucleocapsid assembly. Together, the results demonstrate that the VP24 NES and surrounding residues are important for nucleocapsid assembly, but VP24 nuclear trafficking is not sufficient for the production of infectious viral particles.

While VP24 nuclear trafficking is not sufficient for nucleocapsid assembly, it may still serve an important function in the EBOV life cycle. Previous studies have demonstrated that VP24 can interact with components of the nuclear membrane, such as emerin, lamin B, and lamin A/C [36, 37, 44]. These interactions compromise nuclear membrane integrity, leading to increased activation of signaling pathways involved in nuclear envelope damage [44]. These finding support the possibility that VP24 nuclear transport has biologically significant effects on infected cells. To further elucidate how VP24 nuclear trafficking affects the outcome of infection, future studies should examine how VP24 nuclear trafficking mutations impact nuclear membrane integrity. In addition to nuclear membrane proteins, VP24 interacts with proteins from various cellular compartments [36, 37]. How nuclear trafficking may affect these interactions remains to be determined. Together, the results in this study demonstrate critical assembly functions related to VP24 interactions with the cellular nuclear trafficking machinery.

## Materials and Methods

### Cell culture

Experiments were performed using Human embryonic kidney (HEK293Ts) (ATCC, CRL-3216) and Huh7s (a generous gift from the Gordon lab at the University of California at San Francisco). Cells were maintained in Dulbecco’s Modified Eagle Medium (DMEM). DMEM was supplemented with 10% fetal bovine serum (10%) and 1% penicillin-streptomycin. Cells were cultured in 5% CO_2_ at 37°C.

### Plasmids

The tetracistronic EBOV minigenome was synthesized by Genescript and is based on a previously described tetracistronic minigenome [20]. The synthesized tetracistronic minigenome was cloned into the plasmid pM1. The plasmids pCAGGS-EBOV VP35, - NP, -VP30, -L were kind gifts from Thomas Hoenen and Heinz Feldmann (Rocky Mountain Laboratories, NIAID) and have been described previously [50]. The plasmid pCAGGS-firefly luciferase has been previously described [50]. All mutant VP24 minigenomes and mutant pCAGGS VP24 plasmids were generated by overlapping PCR. The VP24 expression plasmid pCAGGS VP24 was previously described [22]. The pCAGGS-Flag-KPNA4, pCAGGS-Flag-KPNA6 which produce tagged IMPA3 and IMPA7, respectively, have been previously described [23].

### EBOV Tetracistronic Minigenome Assay

For a P0 transfection, Huh7 cells were seeded at a density of 2.0×10^4^ cells per well of 96-well plates. A ratio of 1ug DNA to 3ul TransIT LTI (Mirus Bio, Madison, WI, USA) was used to transfect cells with 20.8ng VP35, 20.8ng NP, 12.5ng VP30, 166.7ng L, 41.6ng T7 RNA polymerase, and 41.6ng tetracistronic EBOV minigenome. Empty pCAGGS vector was used in place of L for the –L control. Additionally, 0.8ng per well of firelfy luciferase plasmid was included as a transfection control. To examine suppression of the tetracistronic minigenome by VP24, 10ng or 20.8ng of the indicated pCAGGS VP24 plasmid were also incorporated into the P0 transfection with empty pCAGGS vector as a control. Each transfection was performed in triplicate. Seventy-two hours post-transfection, supernatants containing trVLPs were collected and the remaining cells were lysed for luciferase assay using the 5x Promega Passive lysis buffer and luciferase activity was measured using a The Dual-Luciferase® Reporter (DLR™) Assay System (Promega, Madison, WI, USA). After confirming that firefly luciferase values remained consistent between conditions, we analyzed the average *Renilla* luciferase activity for each sample. After the luciferase assay, the lysates from the triplicate wells were pooled and used to analyze protein expression by western blot.

For a P1 transfection, Huh7 cells were seeded in a 96-well format at the same density as the P0 transfection described above. Using LT1, cells were transfected with 20.8ng VP35, 20.8ng NP, 12.5ng VP30, 166.7ng L and 0.8ng firefly luciferase plasmid. Each transfection was performed in triplicate. Twenty-four hours post-transfection, cells were infected with 5ul or 50ul of trVLPs generated from a prior P0 transfection. Seventy-two hours post trVLP infection, luciferase activity was measured as described above. Luciferase assay results were determined using a BioTek Cytation C10 Confocal Imagining Reader (Agilent Technologies, Lexington, MA, USA). Lysates from triplicate wells were pooled and analyzed by western blot.

For nuclear and cytoplasmic extractions, a P0 transfection was performed in a 6-well format in which Huh7 cells were seeded at a density of 6×10^5^ cells per well. Using LT1, 125ng VP35, 125ng NP, 75ng VP30, 1000ng L, 250ng T7 RNA polymerase, and 250ng of the tetracistronic EBOV minigenome were transfected. Seventy-two hours post-transfection, NE-PER Nuclear and Cytoplasmic Extraction Reagent (Thermo Fisher Scientific, Waltham, MA, USA) was used to separate the nuclear and cytoplasmic fractions per the manufacturer’s instructions. The nuclear and cytoplasmic fractions were analyzed by western blot.

### RNA Extraction and Quantitative RT-PCR

For RNA extraction, a P0 transfection was performed in a 6-well format. Seventy-hours post-transfection, trVLP supernatants from the P0 transfection were collected and used for a P1 infection. For P1, Huh7 cells were transfected with 125ng VP35, 125ng NP, 75ng VP30, and 1000ng L in a 6-well format. Twenty-four hours post-transfection, P1 wells were infected with 120ul trVLPs from the prior P0 transfection. Seventy-two hours post trVLP infection, transfected P1 cells were lysed with Trizol Reagent (Thermofisher Scientific, Waltham, MA, USA) and RNA was extracted using Direct-zol RNA miniprep kits (Zymo Research, Irvine, CA, USA) per the manufacturer’s instructions. cDNA was generated from 100ng of purified RNA using Random Hexamers and the SuperScript IV First Strand Synthesis System (Thermo Fisher Scientific, Waltham, MA, USA). Quantitative PCR was performed using PerfeCTa SYBR Green SuperMix (Quantbio, Beverly, MA, USA). Quantitative PCR was performed with biological and technical duplicates. GAPDH was used as an endogenous housekeeping gene to calculate delta delta cycle thresholds. Results are represented as fold expression relative to the -L control. Primers targeting VP40 and EBOV 5’Trailer were first described in Galao et al, 2022 [51]. GAPDH forward: CCATGTTCGTCATGGGTGTG. GAPDH reverse: GGTGCTAAGCAGTTGGTGGTG.

### Purifying trVLPs through a sucrose cushion

A P0 transfection in a 6-well format was performed. Seventy-two hours post-transfection, supernatants containing trVLPs were harvested and centrifuged for 5 minutes at 2500 rpm in a 5920R Eppendorf benchtop centrifuge to remove cell debris. Supernatants were then applied to a 20% sucrose cushion in NTE buffer (10mM Tris, 1mM EDTA, 100mM NaCL) and centrifuged at 36,000 rpm in a SW40-TI rotor for 1.5 hours at 4°C in a Beckman L7 ultracentrifuge. trVLPs were then resuspended in NTE buffer and viral protein levels were analyzed by western blot. P0 transfected cells used to generate the trVLPs were lysed in 1% NP-40 buffer (50 mM Tris pH 7.5, 280 mM NaCl, 0.5% NP-40, 0.2 mM EDTA, 2 mM EGTA, 10% glycerol, protease inhibitor (complete; Roche, Indianapolis, IN, USA)) and analyzed by western blot.

### Immunofluorescence assays

Huh7 cells were seeded on coverslips at a density of 1×10^5^ cells per well of a 24-well plate. A ratio of 1ug DNA to 1.5ul lipofectamine2000 (Thermo Fisher Scientific, Waltham, MA, USA) was used to transfect 500ng L, 62.5ng NP, 62.5ng VP35, 37.5 ng VP30, 125 ng T7 RNA polymerase, and 125 ng tetracistronic EBOV minigenome. Empty vector pCAGGS was used in place of L for the –L control. For leptomycin B (LMB) experiments, we performed a P0 transfection as described above and 24 hours post transfection cells were treated with 50nM of LMB. Cells were fixed with 4% paraformaldehyde and permeabilized with 0.1% Triton X-100 seventy-two hours post-transfection. Cells were incubated overnight with rabbit anti-VP24 antibody (SinoBiological, cat #40454-T46) alone or with mouse anti-VP35 monoclonal antibody 6C5 [52]. After incubation with primary antibody, the coverslips were washed three times and incubated with anti-rabbit Alexa Fluor 488 Thermofisher, cat #A-11034) or anti-mouse Alexa Fluor 488 (Thermofisher, cat #A-11001) and anti-rabbit Alex Fluor 647 (Thermofisher, cat #A-11034) secondary antibodies. Nuclei were stained using Hoechst 33342, trihydrocholoride trihydrate (Invitrogen, cat #H3570). To calculate the average nuclear fluorescence over average cytoplasmic fluorescence for VP24, the BioTek Cytation 10 Confocal Imaging Reader (Agilent Technologies, Lexington, MA, USA) was used. Images of the same coverslips were captured using the BioTek Cytation 10 at 60x or using a Leica TCS SP8 at 63x at the Microscopy and Advanced Bioimaging CoRE at the Icahn School of Medicine at Mount Sinai.

### ISRE/IFN-β Reporter Gene Assays

HEK293T cells were seeded at a density of 7.0×10^4^ cells per well in 96-well plates. At a ratio of 1ug DNA to 3ul reagent, TransIT LTI (Mirus Bio) was used to transfect 30ng of either IFNβ promoter or ISRE promoter-firefly luciferase reporters and 30ng pRL-TK *Renilla* luciferase. In addition to the reporter plasmids, 25ng, 50ng, or 75ng of VP24 WT and mutant plasmids were transfected per well. Empty pCAGGS vector was used as a control. Twenty-four hours post transfection cells were treated with either 1000 units/ml of universal type I IFN (PBL Assay Science, NJ, USA) for the ISRE reporter or Sendai virus Cantell strain for the IFN-β reporter. Twenty-four hours post-treatment, luciferase activity was measured as described above. Firefly luciferase values were normalized to *Renilla* luciferase values. Each reaction was performed in triplicate. Triplicate wells were pooled and used to analyze protein expression by western blot.

### Co-immunoprecipitation

HEK293T cells were seeded at a density of 1×10^6^ cells per well in 6-well plates. Using Lipofectamine2000 (Thermofisher Scientific, Waltham, MA, USA) at a ratio of 1ug DNA to 1ul reagent, cells were transfected with 2ug VP24 and 2ug HA-IMPA6 or empty vector pCAGGS. Twenty-four hours post-transfection, cells were lysed in 1% NP40 lysis buffer. Cell lysates were incubated with EZview anti-HA agarose affinity gel (Sigma Aldrich, St. Louis, MO, USA) for 1 hour at 4°C with rocking to precipitate HA-IMPA6. Beads were then washed four times with NP-40 lysis buffer. Bound material was eluted by incubating the beads with HA peptide (Sigma Aldrich, St. Louis, MO, USA) for 30 minutes at 4°C. Whole cell lysates and co-immunoprecipitation samples were analyzed by western blot.

### Western blot

Lysates were run on 4-12% Bis Tris polyacrylamide gels (Thermofisher Scientific, Waltham, MA, USA) and transferred to a PVDF membrane Sigma (Aldrich, St. Louis, MO, USA). Five percent non-fat dry milk in phosphate buffer saline with 0.1% Tween20 (PBST) was used as blocking buffer. Blots were probed with the indicated antibodies and developed using the Western Lighting plus ECL (Perkin Elmer, Waltham, MA, USA). Blots were imaged on the Bio-Rad ChemiDoc MP Imaging System (BioRad, Philadelphia, PA, USA). Antibodies: Rabbit anti-VP24 (SinoBiological, cat #40454-T46); Mouse anti-VP35 (6C5) [52]; Mouse anti-VP40 (IBT Bioservices, cat #0201-016); Rabbit anti-VP35 (IBT Bioservices, cat #0301-040); Rabbit anti-NP (IBT Bioservices, cat #0301-012); Rabbit anti-HA (Invitrogen, cat #71-5500); β-Tubulin (Sigma Aldrich, cat #T8328); Rabbit anti-HDAC2 (Cell Signaling, cat #2540S).

### Electron Microscopy

Huh7 cells were seeded at a density of 2.5×10^5^ cells per well in a 6-well plate. Twenty-four hours after seeding, cells were transfected 1ug NP, 1ug VP35, and 1ug VP24. As a control, empty pCAGGS vectors was used instead of VP24. Cells were transfected with Lipofectamine2000 (Thermofisher Scientific, Waltham, MA, USA) at a ratio of 1ug DNA to 1ul reagent. Twenty-four hours post transfection cells were fixed in 2% glutaraldehyde in 0.1 M sodium cacodylate buffer (pH 7.2) 2 mM CaCl_2_ for over 1 hour at room temperature and post-fixed in 1% osmium/0.8% potassium ferricyanide in 0.1 M cacodylate buffer. This was followed by post-staining in 1% uranyl acetate in water, dehydration in an ethanol series, and embedding in Eponate 12 (Ted Pella, Inc). Ultrathin sections (60–65 nm) were stained with uranyl acetate and lead citrate, and images were acquired using a Tecnai 12 Spirit transmission electron microscope (FEI, Hillsboro, Oregon, USA) operated at 120 kV, equipped with an AMT BioSprint29 digital camera.

### EBOV Infection and Confocal Microscopy

Huh7 cells were seeded at a density of 20,000 cells per well on 8-well Ibidi u-Slides (Ibidi, cat #80826-90). The following day, cells were transfected with Lipofectamine 3000 (Invitrogen, cat #L3000015) with 750ng of either pCAGGS-KPNA4-FLAG (encodes IMPA3), pCAGGS-KPNA6-FLAG (encodes IMPA7), or no plasmid as a transfection control. Media was replaced 18 hours post-transfection and the cells challenged with sucrose-purified EBOV-Mayinga 30 to 40 hours post-transfection at an MOI of 0.3. Cells were then fixed in buffered formalin 20 hours post challenge. This was done for two biological replicates.

Cells were permeabilized with 0.1% Triton X-100 for 10 minutes and blocked in 3.5% BSA for 1 hour. Mouse anti-flag (Sigma-Aldrich, cat #F1804) and rabbit anti-VP24 (Sino Biological, cat #40454-T46) antibodies were added at a 1:2000 dilution and incubated overnight at 4°C. Slides were washed three times with 1x Tris-Buffered Saline with 0.1% Tween (TBST) and incubated with goat anti-mouse AlexaFluor 488 (Invitrogen, cat #A11029) and goat anti-rabbit AlexaFluor 594 Plus (Invitrogen, cat #A32740) at a 1:1000 dilution for 1 hour at room temperature. Cells were washed again three times in 1x TBST and 1:10,000 Hoechst 33342, trihydrochloride trihydrate (Invitrogen, cat #H3570) was added.

Slides were imaged on a Nikon Eclipse Ti2 microscope with AX/NSPARC confocal scan head using a 60x oil lens. Image channel intensities were kept constant and background set using the mock transfected cell controls in ImageJ Fiji software [53].

## Acknowledgments

This work was supported by NIH grants R01AI148663 to CFB and P01AI120943 (Amarasinghe) to CFB and RAD. O.A.V. and was supported in part by Public Health Service institutional research training award AI07647. We thank Emma Komers for expert technical assistance.

## Supporting Information Captions

**S1 Figure:**
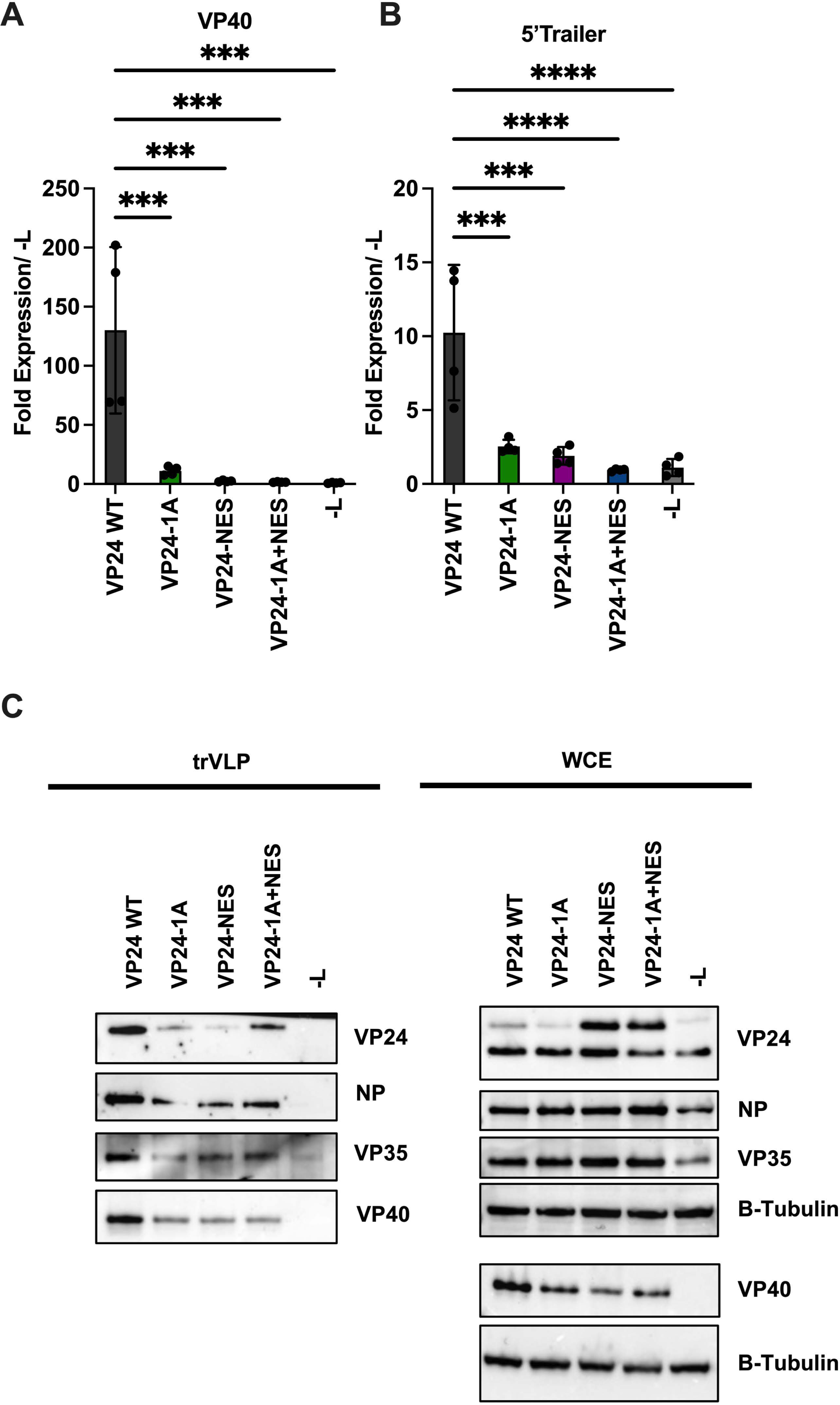
Viral Genome Replication and trVLP Production are Reduced for VP24 Mutant trVLPs. A-B) Quantification of viral RNA at P1. P0 trVLP supernatants generated from the indicated minigenomes were harvested and used to infect transfected target cells at P1. –L indicates a P0 transfection with the VP24 WT minigenome in which the L expression plasmid was omitted. Seventy-two hours post-infection, cells were harvested for RNA extraction and cDNA was generated using random hexamer as primer. Reverse transcription-quantitative PCR (RT-qPCR) was performed using primers targeting VP40 (A) or the EBOV 5’ trailer (B), as described previously [51]. Data are represented as fold expression relative to the -L control. **** denotes p-value ≤ 0.001. *** denotes p-value ≤ 0.001. C) Western blot analysis of viral protein expression in trVLPs from P0. Following a P0 transfection in Huh7 cells, trVLP supernatants were concentrated by ultracentrifugation through a sucrose cushion. The concentrated trVLPs were analyzed by western blot for viral protein expression. The remaining P0 transfected cells were lysed and the whole cell extracts (WCE) were analyzed by Western blot for viral protein expression.

**S2 Figure:**
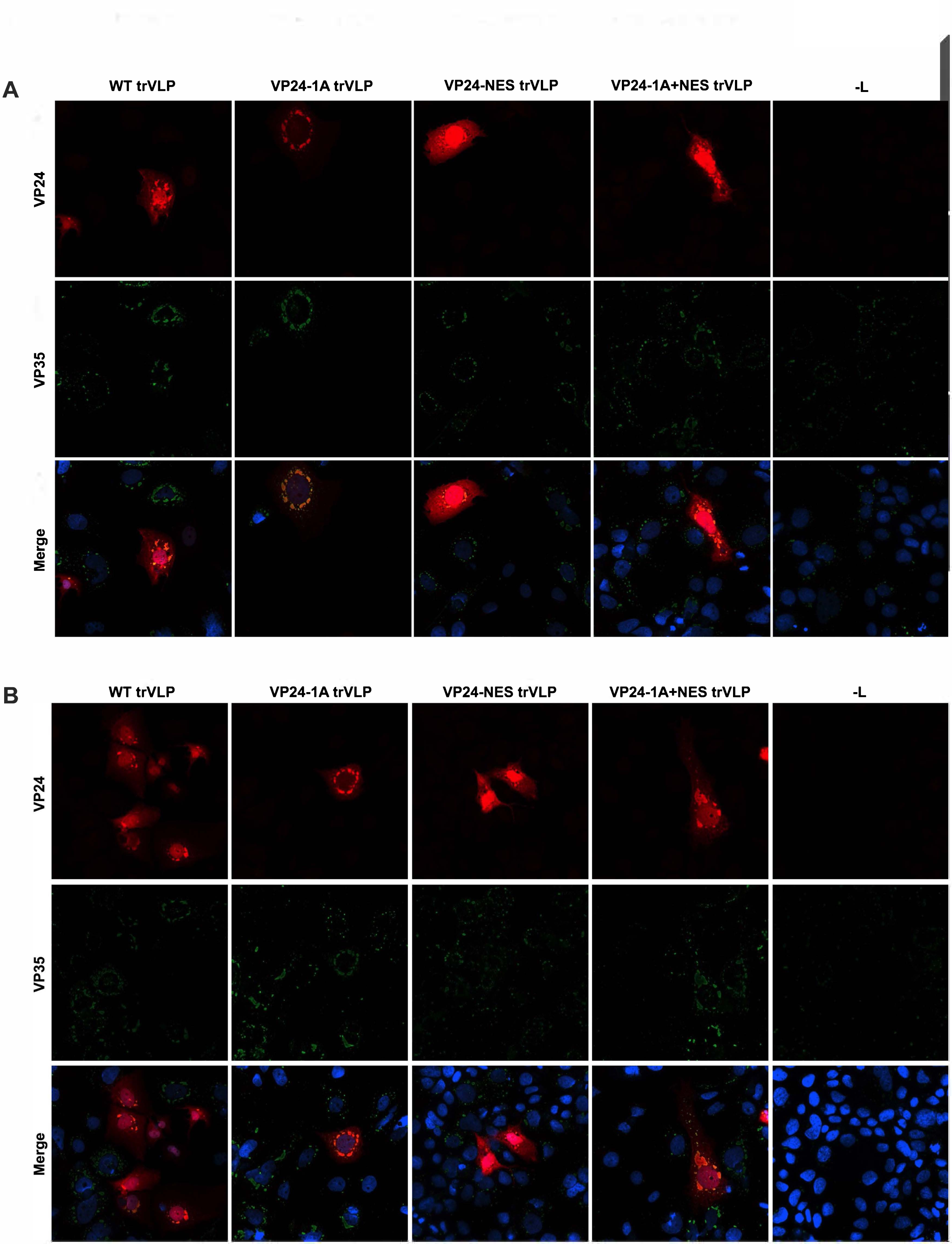
VP24-WT, VP24-1A, VP24-NES, and VP24-1A+NES Colocalize with Viral Inclusion Bodies. A-B) Confocal microscopy images of P0 transfected Huh7 cells. Seventy-two hours post-transfection with the indicated minigenomes, cells were fixed and stained for VP24, VP35, and nuclei. Confocal microscopy images were captured using a Biotek Cytation C10 at 60x.

**S3 Figure:**
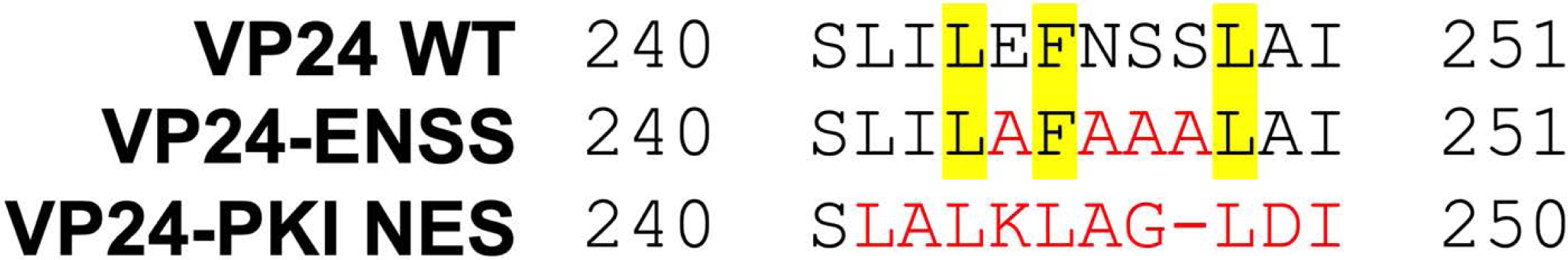
Alignment of VP24 WT, VP24-ENSS, and VP24-PKI NES C-termini. An alignment of the 240-251 C-terminal residues for the VP24 WT, VP24-ENSS, and VP24-PKI NES. Yellow highlighted residues indicate the VP24 WT NES. Residues in red indicate residues that differ from the VP24 WT sequence. The hyphen was inserted into the PKI NES to align the C-terminal most L and I with those in the other sequences.

